# Type 1 Diabetes: an Association Between Autoimmunity the Dynamics of Gut Amyloid-producing *E. coli* and Their Phages

**DOI:** 10.1101/433110

**Authors:** George Tetz, Stuart M. Brown, Yuhan Hao, Victor Tetz

**Author notes:** Correspondence and requests for materials should be addressed to G.T.

## Abstract

The etiopathogenesis of type 1 diabetes (T1D), a common autoimmune disorder, is not completely understood. Recent studies suggested the gut microbiome plays a role in T1D. We have used public longitudinal microbiome data from T1D patients to analyze amyloid-producing bacterial composition and found a significant association between initially high amyloid-producing *Escherichia coli* abundance, subsequent *E. coli* depletion prior to seroconversion, and T1D development. In children who presented seroconversion or developed T1D, we observed an increase in the *E. coli* phage/*E. coli* ratio prior to *E. coli* depletion, suggesting that the decrease in *E. coli* was due to prophage activation. Evaluation of the role of phages in amyloid release from *E. coli* biofilms *in vitro* suggested an indirect role of the bacterial phages in the modulation of host immunity.

This study for the first time suggests that amyloid-producing *E. coli*, their phages, and bacteria-derived amyloid might be involved in pro-diabetic pathway activation in children at risk for T1D.

## Introduction

Type 1 diabetes (T1D) is an autoimmune disorder driven by T cell-mediated destruction of the insulin-secreting β-cells of the pancreatic islets that often manifests during childhood (Lehuen et al., 2010). There are a number of factors associated with the development of T1D (Atkinson and Eisenbarth, 2001; Soltesz et al., 2007; Rewers, M., and Ludvigsson, 2016). The strongest susceptibility alleles for T1D encode certain human leukocyte antigens (HLAs), which play a central role in autoreactive T-cell activation in T1D pathogenesis (Barrett et al., 2009; Ikegami et al., 2011). Only a portion (3–10%) of children carrying HLA risk alleles will develop T1D, pointing out the role of environmental factors in disease initiation; however, the role of environmental factors in T1D etiology remains elusive (Achenbach et al., 2005; Noble and Valdes, 2007; Rewers, M., and Ludvigsson, 2016).

T1D is generally associated with a long pre-diabetic seroconversion period, during which autoantibodies to antigens of pancreatic β-cells or insulin are produced (Vehik et al., 2011). There are a few known factors that drive autoimmunity during infancy, such as so-called “natural apoptosis” or spontaneous cell death within the β-cell population, deposition of islet amyloid polypeptide (IAPP) aggregates, microbiota alterations, or viral infection that specifically targets pancreatic β-cells and leads to islet cell death, which contribute to the formation of β-cell antigens, activation of dendritic cells (DC), and antigen presentation (Drescher et al., 2004; Vaarala et al., 2008; Eizirik et al., 2009; Lehuen et al., 2010; Todd, 2010, Paulsson et al., 2014; Knip and Siljander, 2016). However, the exact roles of these mechanisms in disease onset remain poorly understood (Lehuen et al., 2010).

Pancreatotropic viruses, such as enterovirus, coxsackie B, mumps, rubella, and cytomegalovirus, have been evaluated as triggers of T1D since the discovery of antiviral antibodies in diabetic patients and because of the role of viral infection in elevated levels of type I interferon (IFN), which is known to play a role in T1D progression (Flodström et al., 2002; Op de Beeck and Eizirik 2016; Hyöty 2016). However, in many patients who develop T1D, there is no clear link with viral infections (Filippi and von Herrath, 2008). Moreover, the mimicry between β-cell autoimmune antibodies and viral epitopes of coxakieviruses, which are the most strongly associated with T1D, has recently been shown not to underlie T1D pathogenesis (Schloot et al., 2001; Harrison 2019). Natural apoptosis has been described only in certain patients and highlights the gaps in our knowledge about disease initiation (Lehuen et al., 2010;). Therefore, despite decades of research, the etiology of autoimmune response induction remains elusive (Harrison, 2019).

Previous studies have established a correlation between seroconversion and chronic inflammation resulting from impaired gut barrier function and increased intestinal permeability, which is suggested as a hallmark of T1D (Vaarala et al., 2008). Given the overarching influence of gut bacteria on human health, including intestinal permeability and its regulatory role in autoimmunity, the role of the microbiota in T1D development has been recently explored (Knip and Siljander, 2016; Kostic et al., 2016; Paun et al., 2017). The microbiota of the human intestinal tract is comprised of bacteria, fungi, and eukaryotic and bacterial viruses (bacteriophages). Bacteria in the human gut live within surface-associated microbial communities termed biofilms, which are characterized by the presence of self-produced extracellular matrix and a surface film that protects the microorganisms from the outer environment (Costerton et al., 1999; Tetz et al., 2009; Flemming and Wingender, 2010; O’Toole et al., 1999). The extracellular matrix consists of different biomolecules, including extracellular nucleic acids, polysaccharides, and proteins. Several microorganisms within the human microbiome, predominantly members of *Enterobacteriaceae*, also produce amyloid proteins that can form so-called “curli fibers” which provide biofilms with unique mechanical properties (Chapman, 2002; Barnhart and Chapman 2006). Bacterial curli proteins of *Enterobacteriaceae* form highly ordered β-sheet rich amyloid composed of a major subunit, CsgA, and a minor subunit, CsgB (Schwartz and Boles 2013). Bacterial amyloids share certain similarity with human amyloids; however, whereas amyloid in bacteria is suggested to be physiological, in mammals, the formation of misfolded, insoluble, highly ordered prion-like cross-β structures is commonly associated with a variety of diseases (Fowler et al., 2007). In humans, pathological deposition of insoluble amyloid aggregates has been shown to be associated with the development of Creutzfeldt-Jakob disease, Alzheimer’s disease, Parkinson’s disease, and T1D, where an increased islet amyloid polypeptide (IAPP) concentration may constitute a risk factor for β-cell destruction (Paulsson et al., 2014; Collinge 2016). Moreover, similarities between bacterial and eukaryotic amyloid β-sheets have been implicated in inter-kingdom communication through the toll-like receptor signaling pathway, leading to the deposition of misfolded protein in the brain and altered amyloidogenesis in *Caenorhabditis elegans* and rats following colonization with amyloid-producing *E. coli* (Chen et al., 2016). Notably, in this work, the authors showed that bacterial curli fibers triggered T-cell immunity through the TLR-2-MyD88-NF-kB signaling pathway, which is also known to be associated with T1D progression; however, no microbiome studies have been conducted to reveal the association of amyloid-producing bacteria with diabetes (Tukel et al., 2010). Moreover, only a limited number of studies on the human microbiome have revealed alterations that may be associated with T1D. Kostic et al. and Zhao et al. found that an increased abundance of *Blautia* spp., *Ruminococcus* spp., and OTUs belonging to the genus Bacteroides positively correlated with T1D progression, suggesting that the primary pathological implication in autoimmunity might be through an altered metabolite profile associated with these bacteria, including triglyceride and branched-chain amino-acid production Davis-Richardson et al. 2014; Kostic et al., 2016). Notably, a previous microbiota study revealed microbiome alterations that emerged only after the appearance of autoantibodies, suggesting that such alterations are involved in clinical progression to T1D rather than in disease initiation (Knip, and Siljander 2016).

There are multiple causes of gut bacterial alterations in T1D, and bacteriophages are considered to be one of the most influential, but are poorly studied in this regard (Zhao et al., 2017). Data from a phagobiota analysis of children with T1D are even more limited, and showed that alterations in Bacteroides phages are discriminative for disease status, but not clearly associated with disease progression (Zhao et al., 2017). Bacteriophages as possible agents that may negatively affect mammalian health have attracted scientific attention only recently (Tetz and Tetz 2018). We have previously shown that bacteriophages can alter the mammalian microbiota, leading to increased intestinal permeability and triggering chronic inflammation, which potentially participate in protein misfolding and might contribute to the onset of Parkinson’s disease Tetz et al., 2017; Tetz et al., 2018).

In this study, we conducted a detailed comparative metagenomic analysis of amyloid-producing gut bacteria and the associated phagobiome in a prospective, longitudinal cohort of HLA-matched infants up to 3 years of age. We retrieved a dataset of short sequence reads generated by Kostic et al. (2015) from the NCBI Sequence Read Archive (SRA) (https://www.ncbi.nlm.nih.gov/bioproject/382085) and analyzed amyloid-producing bacteria and bacteriophage diversity using MetaPhlAn and a custom method (Segata et al., 2012; Kostic et al., 2016).

This study for the first time revealed alterations in the abundance of intestinal amyloid-producing bacteria during early life due to prophage activation that that correlated with the development of T1D-associated autoimmunity. These data were supported by findings *in vitro* that revealed significant amyloid release from *E. coli* biofilms upon induction by prophages, suggesting a potential involvement of *E. coli*-derived curli fibers in pro-diabetic activation pathways in children at risk for T1D.

## Results

### Autoimmunity Development is Associated with an Alteration in Amyloid-producing Bacterial Abundance

To explore the potential link between the abundance of amyloid-producing bacteria and T1D-associated seroconversion, we used longitudinal shotgun metagenomics sequencing data of the fecal microbiome from 10 children who exhibited autoantibodies (six seroconverters and four children who developed T1D) and eight non-seroconverted HLA-matched control individuals. All patient characteristics were previously reported by Kostic et al. (Table S1) (Kostic et al., 2016). HiSeq-2500 sequencing was used, producing an average of ~2.5 Gb per sample. Amyloid-producing bacteria were represented by *E. coli*, *Staphylococcus aureus*, and *Salmonella spp.* (Barnhart and Chapman 2006; Schwartz and Boles 2013). Among these amyloid-producing bacteria, *E. coli* was the major group, while the other curli-producing bacteria were identified only in a single sample and at a few collection times and thus, were disregarded in subsequent analysis (Table S2). We first compared the association between *E. coli* abundance, disease phenotype, and collection time using a two-tailed Mann–Whitney *U* test (Figure 1; Table S3). We clustered the samples into time-point bins of 300 days, and studied the dynamics of *E. coli* abundance over 1300 days within and among patient groups. *E. coli* abundances demonstrated a different dynamic relationship across time for case and control groups. *E. coli* tended to disappear in T1D and seroconverters over the time, whereas in controls, *E. coli* abundance tended to increase and did not change significantly over time. In both the seroconverter and T1D groups, we found a statistically significant decrease in *E. coli* abundance when comparing the first 0-300 day period and the following 300-600, 600-900, 900-1300 time bins (*p* < 0.05). Notably, the initial abundance of *E. coli* was significantly higher in cases than in controls at 0–300 days. The case groups responded differently over time, and importantly, data from the control group revealed that there was no variation in *E. coli* associated with age.

**Figure 1.**
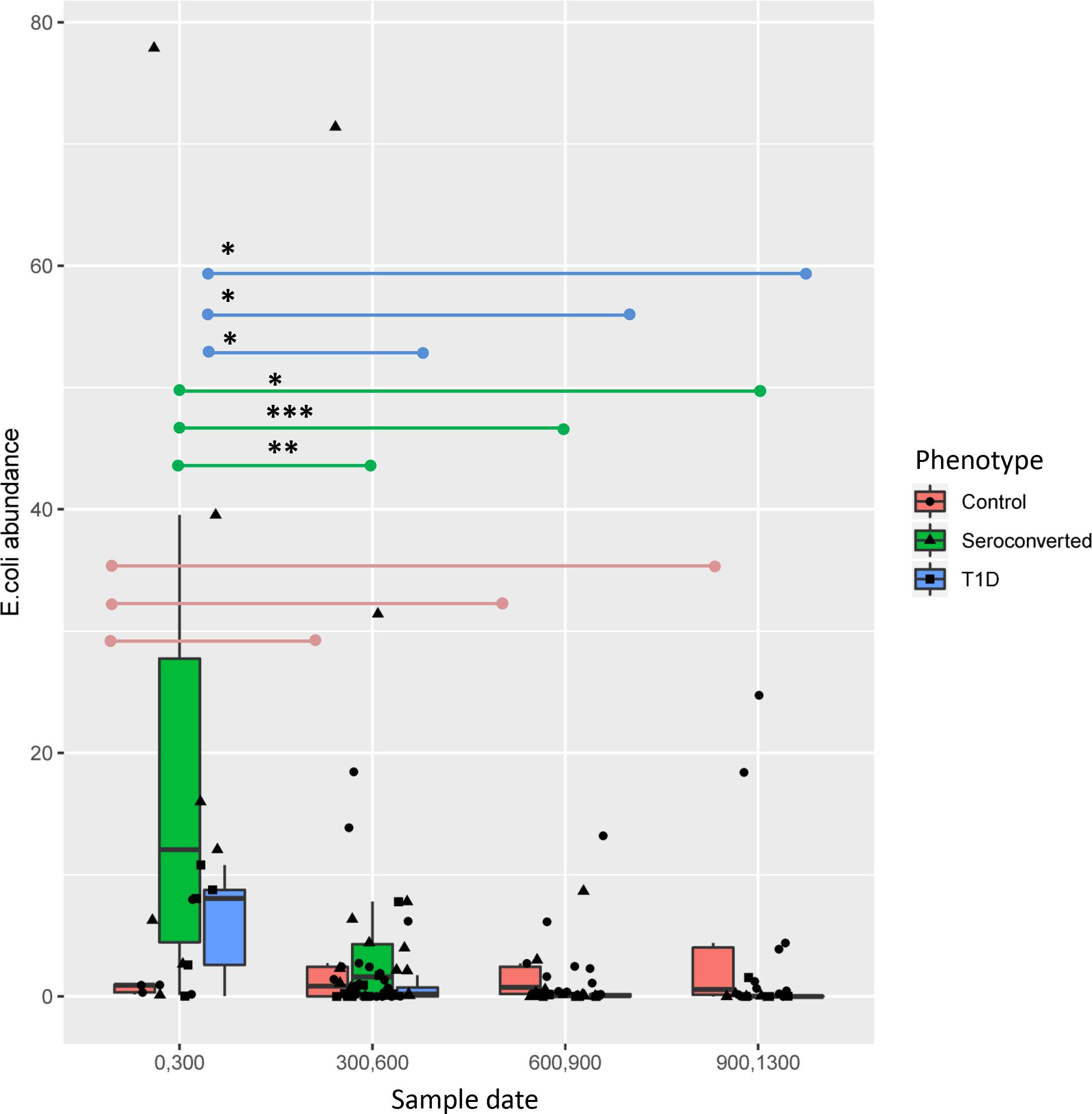
*E. coli* Abundance Over Time. Boxplot showing absolute *E. coli* abundances in the seroconverter, T1D, and control groups in the indicated periods. *E. coli* abundance across different time points was compared using two-tailed Mann-Whitney U test. Statistically significant variation between selected conditions (*p < 0.05, **p < 0.01 and*** p < 0.005).

To explore the relation between *E. coli* abundance and autoimmunity further, we next analyzed whether alterations in the abundance of *E. coli* could be associated with the development of autoantibodies in more detail, and whether or not *E. coli* abundance could distinguish the T1D disease state. To this end, we studied the appearance and disappearance of *E. coli* in each patient in dynamic association with the appearance of autoantibodies (Figure 2). We found a positive correlation between the total disappearance or presumed disappearance (defined as a >50-fold decrease) in *E. coli* and autoantibody appearance in the T1D and seroconverter groups, but not in controls (Walterspiel et al., 1992). All T1D patients showed disappearance or an episode of disappearance of *E. coli* before the detection of autoantibodies and disappearance of *E. coli* prior to the diagnosis of diabetes (Table S4). In seroconverters, *E. coli* disappeared in all patients, except E003989 and T013815. For patient E10629, data were difficult to interpret because of the long period between sample collections (556 days between the last sample taken before seroconversion and the first sample after seroconversion). In any case, *E. coli* was absent in the first sample following seroconversion; thus, it is reasonable to assume that it had disappeared before autoantibody appearance. Notably, in certain control patients, *E. coli* disappeared at some time points; however, this was followed by a restoration of the *E. coli* population in all cases, which was not regularly seen in case subjects.

**Figure 2.**
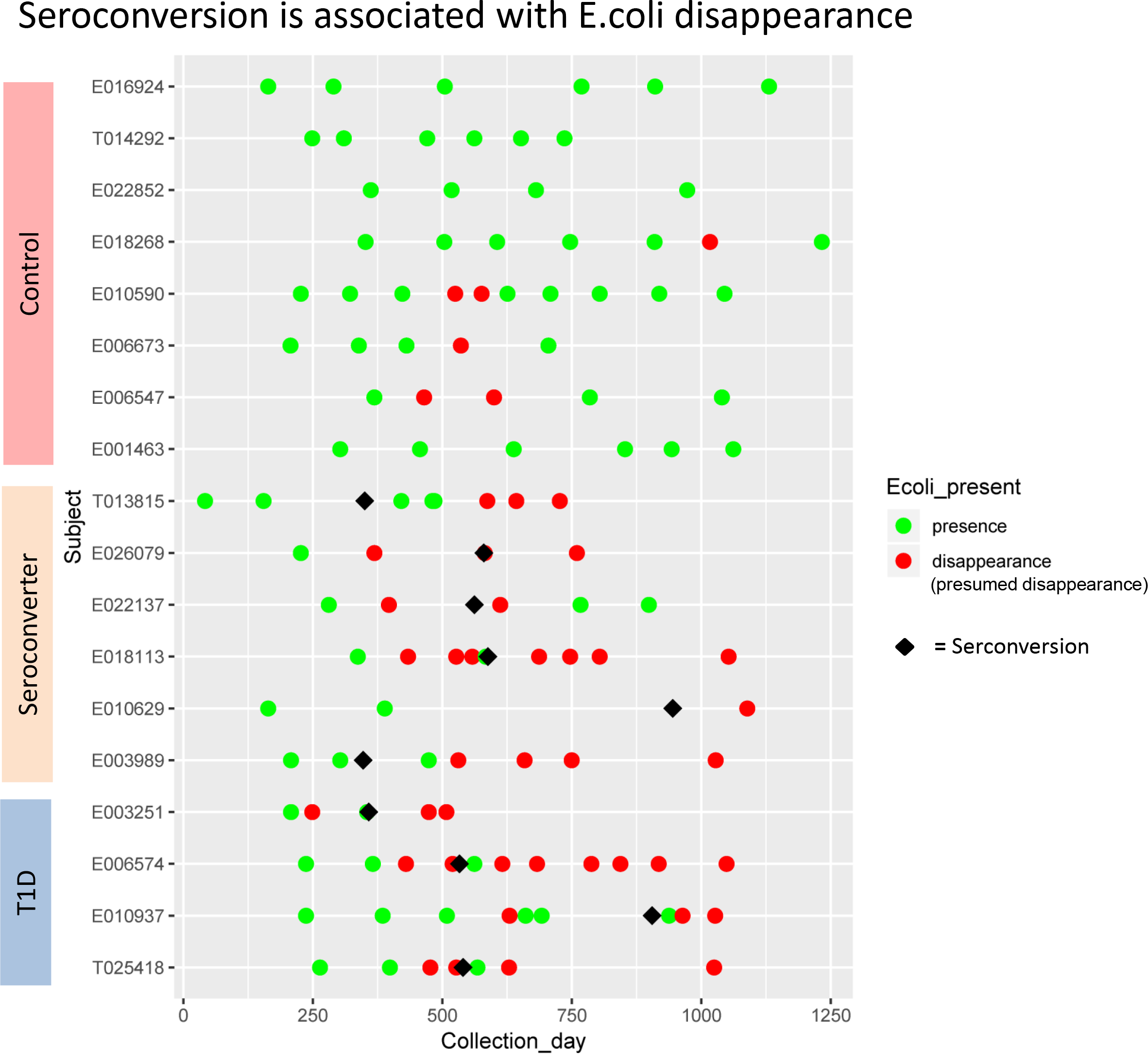
Association Between the Dynamics of the Disappearance of *E. coli* and Seroconversion. Each row represents an individual, with each symbol indicating a single stool sample. The X-axis indicates age at sample collection. *E. coli* abundance is >0.01. Green indicates presence of *E. coli*, and red indicates disappearance or presumed disappearance (over 50-fold reduction when compared with initial abundance) of *E. coli*. Diamonds represent the time point at which autoantibodies were detected.

Next, we examined changes in the absolute abundance of *E. coli* in each patient individually among different periods, and we presented the data as a heatmap (Figure 3). We detected alterations in *E. coli* abundance within children from the control group, followed by restoration of the *E. coli* population to the levels equal to or above the pre-seroconversion levels, whereas restoration of the *E. coli* population to preceding levels was noted in only a few children from the case groups. This more detailed analysis has also revealed some previously overlooked data, for patient T013815. Although in this subject, *E. coli* did not disappear prior to the seroconversion, the appearance of autoantibodies occurred in the period when *E. coli* started to significantly decrease from the highly elevated abundance compared to other patients. Another observation that attracted our attention was that the initial abundance of *E. coli* in T1D and seroconverter groups was significantly higher than that in the control group, which confirmed the data presented in Figure 1. The result further supports that the initial higher abundance of *E. coli* and the decrease of amyloid-producing bacteria abundance in case group were signatures associated with autoimmunity and disease progression.

**Figure 3.**
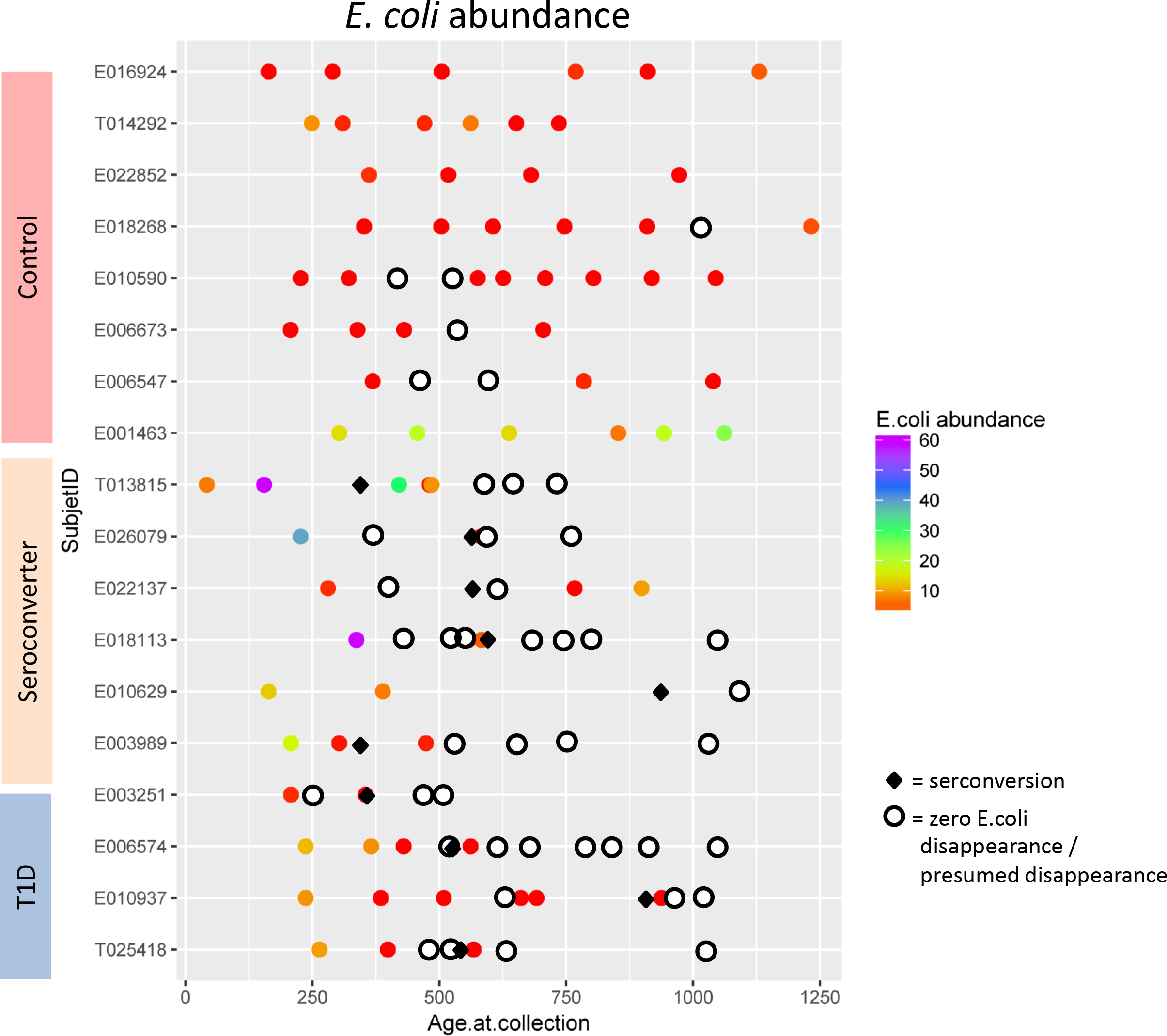
Absolute Abundances of *E. coli* Across Samples. Fecal bacterial communities were analyzed by high-throughput Illumina Hiseq2500 sequencing. Threshold for *E. coli* abundance is >0.01. Each row represents an individual, with *E. coli* abundance in each stool sample (indicated with different symbols) being color-coded using a color gradient, ranging from red (low abundance) to purple (maximum abundance). Samples with no *E. coli* are marked with white circles. The X-axis indicates age at sample collection. Diamond sign represents the time of seroconversion detected.

In addition, we studied the potential correlation between the disappearance of *E. coli* and certain HLA alleles or the appearance of particular autoantibodies, IAA, GADA, IA2A, ZNT8A or ICA, in case groups. No correlations were observed (data not shown). We also detected no association between *E. coli* disappearance and breastfeeding duration, *Bifidobacterium* abundance, or antibiotic usage, which are known to influence *Enterobacteriaceae* (Tables S5, S6) (Candela et al., 2008).

### *E. coli* Bacteriophages as Signatures Associated with Seroconversion

As we identified *E. coli* as T1D-discriminative bacteria, we next studied the relationships between this microorganism and bacteriophages of *E. coli*, as phages are known as main regulators of bacterial populations. First, we detected 63 *E. coli* phage species in case and control groups. We evaluated possible differences in *E. coli* phages, using α-diversity, in the pre-seroconversion period (Table S7). For the samples from the control group, we set the medium time to seroconversion in case groups (540 days) as an artificial benchmark. Phage richness was statistically different between control and seroconversion individuals as indicated by ACE and Chao1 indexes (ACE: *p* = 0.0181; Chao1: *p* = 0.0137), but was not statistically significantly different between control and T1D, most likely because of the small T1D patient cohort (ACE: *p* = 0.6086; Chao1: *p* = 0.6359). *E. coli* phage diversity tended to be lower in the control subjects than in seroconverters (Shannon, *p* = 0.1526; Simpson, *p* = 0.5437), and was significantly lower in controls vs. T1D cases (Shannon, *p* = 0.0055; Simpson, *p* = 0.0248) (Mann–Whitney test) (Table S7).

We observed that nearly all *E. coli* phages were lysogenic, whereas strictly virulent lytic phages, such as *Enterobacteria* phage IME10 or *Enterobacteria* phage 9g, were found only in a few samples (Table S2). The predominance of lysogenic *E. coli* phages clarifies why α-diversity indices revealed trends of decreased evenness and diversity of phages in case groups compared to controls in pre-seroconversion samples, which most likely reflected lower *E. coli* abundance in controls (Table S2).

Next, we analyzed the *E. coli* phage/*E. coli* bacterial cell ratio, which represents the “lytic potential” (Figure 4; Table S8; Table S9) (Waller et al., 2014). To this end, we first normalized the *E. coli* phage abundance to that of *E. coli*. In theory, the phage/bacteria ratio reflects whether or not a prophage is stably integrated within the host bacterial genome (Tetz et al., 2018). A low ratio indicates that prophage is most likely absent in the genomes of part of the bacterial host population, whereas a high ratio suggests active, productive phage-induced bacterial lysis. The phage/bacteria ratio increased in subjects from all groups prior to the decrease in *E. coli* abundance, indicating that productive phage infection was the cause of *E. coli* depletion. Notably, that observation reflected a statistically significant elevation in the phage/bacteria ratio before the appearance of autoantibodies in the T1D and seroconverter groups (Table S10).

**Figure 4.**
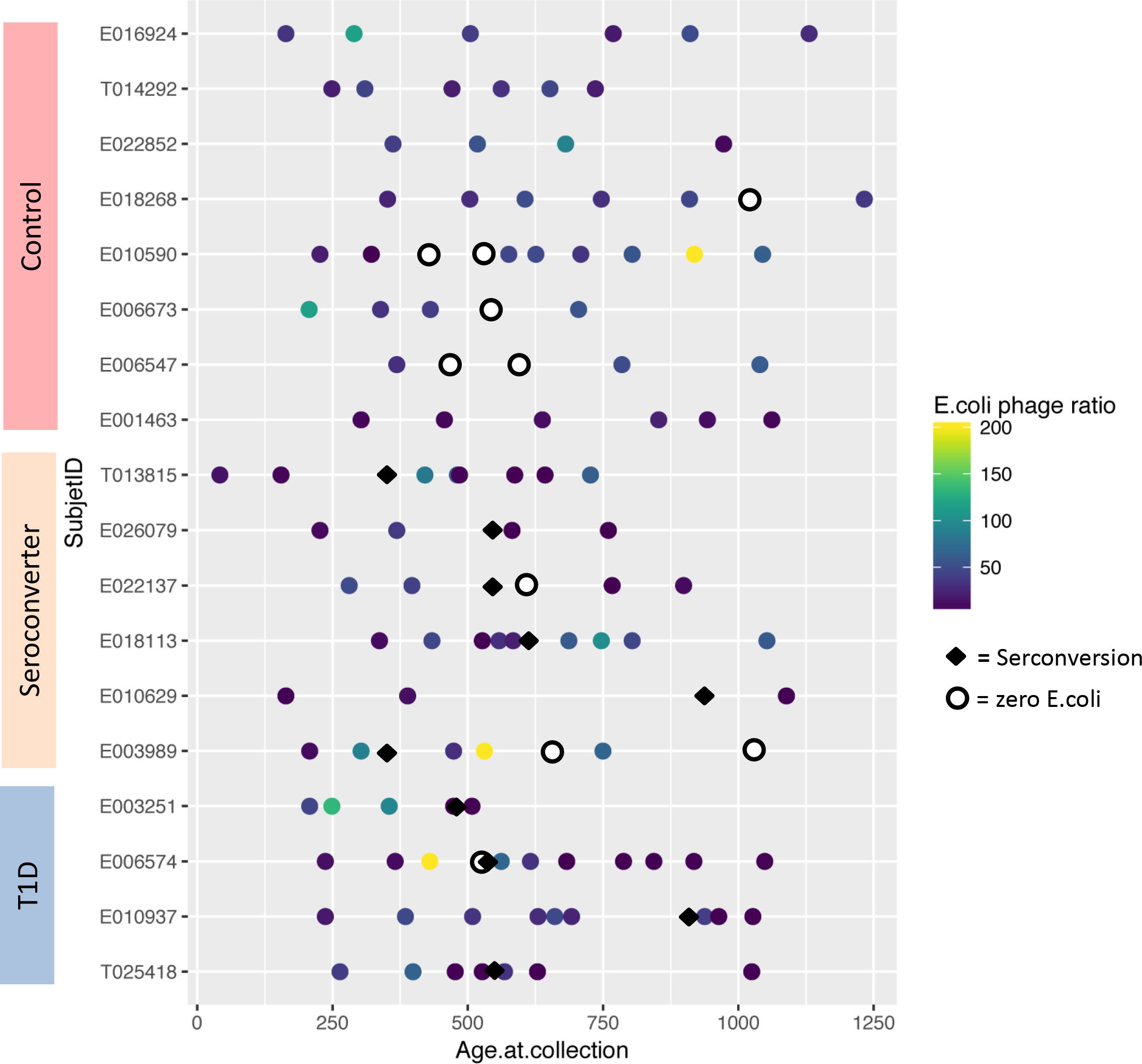
Association Between *E. coli* phage/*E. coli* Ratio Across Samples and Seroconversion. Each row represents an individual, with each symbol representing the phage/bacterial ratio, calculated as *E. coli* phage abundance normalized to that of the *E. coli* abundance, of a sample. Abundance is represented as a color gradient, ranging from purple (lowest ratio) to yellow (maximum ratio). Samples in which *E. coli* phages but no bacterial cells were present are marked with white circles. Samples with total *E. coli* disappearance or presumed disappearance with over 50-fold decrease in *E. coli* are indicated by red circles. The X-axis indicates age at sample collection. The diamond symbol represents the time point at which seroconversion was detected.

To evaluate the implication of the increase in the phage/bacteria ratio as a driving force behind *E. coli* depletion further, we analyzed which phages had a correlation with the depletion of *E.coli* abundance across most subjects. The most frequently found lysogenic phages with an inverse relationship with *E.coli* abundance are the representatives of the *Peduovirinae* subfamily and of the unclassified *Lambdavirus* subfamily within the *Caudovirales* order. The increase in number of the lysogenic E. coli phages along with the decrease in E. coli abundance indicated that there was a productive bacteriophage infection that led to bacterial host death and the release of phage progeny (Waller et al., 2014). These data revealed previously overlooked particularities of the phagobiota in T1D, suggesting a primary role of induction of certain *E. coli* prophages in *E. coli* depletion, and an association with autoimmunity and T1D development.

### Role of Prophage Induction in the Release of Amyloid from *E. coli* Biofilms

We evaluated how the die-off of *E. coli* populations due to prophage induction could lead to amyloid release. We used 48-h old *E. coli* biofilms with confirmed curli expression (curli formation on 48-h colonies was visible with the naked eye on petri dishes (data not shown). Prophages were induced with mitomycin C. Lytic bacteriophage development was confirmed as an increase in plaque forming units (PFU) and a reduction in colony-forming units (CFU). The presence of phage in *E. coli* biofilm was confirmed by the production of plaques on a non-induced *E. coli* control culture between 4 h and 10 h. The effect of prophage induction in *E. coli* biofilm on CFU and PFU is summarized in Table 1. Phages started to appear in the biofilm as of 4 h after prophage induction, with maximum PFU at 8 h, which coincided with a decrease in CFU.

**Table 1.** Effect of prophage induction in MG1655 wild-type lysogenic strain on PFU and CFU.

|  |  | PFU/ml |  |  |  | CFU/ml |  |  |  |
| --- | --- | --- | --- | --- | --- | --- | --- | --- | --- |
|  |  | 4 h | 6 h | 8 h | 10 h | 4 h | 6 h | 8 h | 10 h |
| *E. coli* | control | 0 | 0 | 0 | 0 | 9.3<br>log10<br>+/- 0.6<br>log10 | 8.9<br>log10<br>+/- 0.5<br>log10 | 9.1<br>log10<br>+/- 0.6<br>log10 | 9.5<br>log10<br>+/- 0.6<br>log10 |
| *E. coli* | + mytomicin C | 2.3 log10<br>+/- 0.2<br>log10 | 9.3<br>log10<br>+/- 0.5<br>log10 | 10.4<br>log10<br>+/- 0.6<br>log10 | 11.2<br>log10<br>+/- 0.5<br>log10 | 8.8<br>log10<br>+/- 0.4<br>log10 | 8.2<br>log10<br>+/- 0.7<br>log10 | 7.6<br>log10<br>+/- 0.7<br>log10 | 7.4<br>log10<br>+/- 0.7<br>log10 |

Next, we evaluated the effect of prophage induction on the amount of amyloid fibers released into the biofilm supernatant. To quantify the amount of aggregated amyloid, we evaluated its binding to the amyloid-diagnostic dye Congo red (CR) (Reichhardt et al., 2015). We observed a significant increase in CR absorbance upon binding to supernatant 8 h following prophage induction as compared with control samples (Figure 5). Like for other curli fibers, heating of the *E. coli* supernatant induced a spectral shift of the CR solution, with a maximum difference in absorbance between CR alone and CR bound to amyloid fibers at ~541 nm (Chapman et al.,2002). These data suggested that the phage-mediated lysis of *E. coli* led to the release of amyloid aggregates into the biofilm.

**Figure 5.**
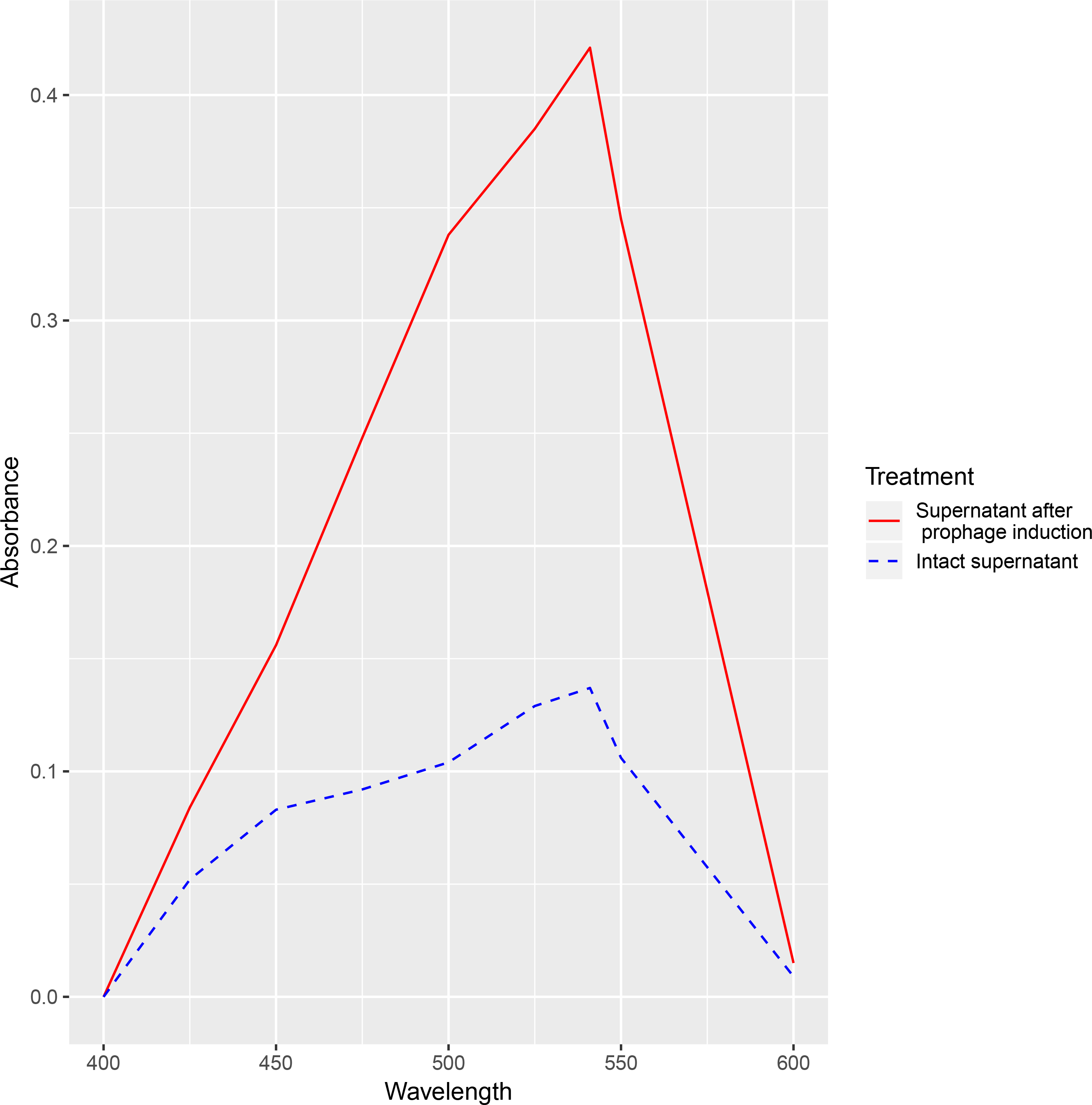
Prophage Induction Promotes Amyloid Release from *E. coli in vitro*. CR absorbance upon binding to amyloid from intact *E. coli* supernatant (dashed line) and from *E. coli* supernatant after prophage induction (solid line). Both curves are approximately equal to zero at the isosbestic point (403 nm), and the maximum difference in absorbance was observed at 541 nm. Both spectra were normalized to the absorbance spectrum of free CR.

## Discussion

This study revealed an association between the dynamics of intestinal amyloid-producing *E. coli* prior to seroconversion in HLA-matched children and the development of T1D-associated autoimmunity. Several lines of evidence support roles of *E. coli*, *E. coli* phages, and *E. coli*-derived amyloid in T1D.

First, in seroconverter and T1D patients, we observed a statistically significant elevation in *E. coli* abundance from day 0 to 300, followed by *E. coli* depletion . Notably, we observed the total disappearance or a pronounced depletion of *E. coli* prior to the appearance of autoantibodies in all children who progressed to T1D. A similar trend was noticed among most patients in the seroconverter group. Based on comparison with the control group, we confirmed that the observed changes in amyloid-producing bacteria were associated with disease state, not with age. Despite the relatively small patient cohort studied, the longitudinal analysis and the large total number of samples (120) analyzed provided sufficient statistical power to identify significant changes (Zhao et al., 2017). There are multiple factors that could lead to *E. coli* depletion within the gut microbiome, including interplay with other gut microorganisms, which constantly change during infancy. *E. coli* is a member of *Enterobacteriaceae*, which are known to positively correlate with *Bifidobacteriaceae*, which are affected by breastfeeding (Candela et al., 2008; Kostic et al., 2016). With that in mind, we evaluated a possible association between these families; however, we found no association between *E. coli*, duration of the breastfeeding, and *Bifidobacteriaceae* abundance. Another environmental factor that can affect the intestinal *E. coli* population is antibiotic usage. Although some children in this study had been exposed to antibiotics, no association between the timing of antibiotic usage and E. coli depletion was noticed (Mills et al., 2013). However, the most profound regulators of bacterial populations are bacteriophages. Notably, we found a positive correlation between the marked decline in *E. coli* abundance and an increase in the *E. coli* phage/*E. coli* bacterial cell ratio. The strongest association was observed for lysogenic *E. coli* phages from the *Peduovirinae* subfamily in the family *Myoviridae* and the number of unclassified *Lambdavirus*, in the family *Siphoviridae* in the order *Caudovirales* suggesting these phages might play a role as a diabetogenic indirectly leading to the release of *E. coli* amyloid-DNA complexes. Most likely, prophage induction that led to *E. coli* depletion was triggered by an external stimulus (Penadés et al., 2015). Prophage induction signals vary among bacteriophages, but generally are activated in response to environmental and internal factors, such as bacterial SOS response molecules triggered by antimicrobial agents, oxidative stress, and DNA damage (De Paepe et al., 2014; Tetz and Tetz, 2018). However, we found no relation between the timing of antibiotic usage and E. coli disappearance relevant to prophage induction.

Significant die-off of the host population by prophage activation results in microbial biofilm disruption, leading to the release of pathogen-associated molecular patterns (PAMPs) (Marrack et al., 200). Accordingly, the *E. coli* die-off observed in this study might have led to the release of a variety of PAMPs, such as LPS, DNA, and amyloid; however, for the majority of PAMPs, their role in T1D remains elusive. Moreover, LPS derived from *E. coli* possess anti-diabetic activity, inhibit innate immune signaling, and enhance endotoxin tolerance in non-obese diabetic mice (Vatanen et al., 2016). The predominant component expressed in *E. coli* biofilms that is unique to amyloid-producing Enterobacteria is curli amyloid fiber, which forms highly immunogenic complexes with DNA (Tursi et al., 2017). Amyloid-DNA composites are known to intensively stimulate TLR2 and TLR9, inducing immunogenic reactions, and they have been associated with the triggering of other autoimmune disorders, including systemic lupus erythematosus (Gallo et al., 2015; Tursi et al., 2017). Therefore, it is reasonable to speculate that die-off of the intestinal *E. coli* population, particularly considering its initial high abundance among children from the case groups, might lead to the prominent release of curli fibers and DNA-amyloid complexes that in turn act as pro-diabetic protein and trigger the autoimmune cascade in susceptible hosts. In favor of the suggested diabetogenic role of *E. coli* amyloid, it has been previously implicated in autoimmunity associated with increased type I IFN-induced activation and protein misfolding (Di Domizio et al., 2012; Gallucci et al., 2015). Type I IFN is a signature cytokine in T1D patients and is known to be one of the injury agents during viral infection, which actually led to the hypothesis of the viral nature of pancreatic damage in T1D (Schloot et al., 2001). Type I IFN is implicated in early T1D stages by triggering β-cell apoptosis and the formation of β-cell autoantigens (Op de Beeck and Eizirik 2016). Notably, *E. coli* curli-DNA composites have the potency to stimulate the type I IFN response via two ways: through TLR stimulation of β-cells and of DCs (Gallo et al., 2015). Indeed one of the pathways underlying the potential diabetogenic action of *E. coli* amyloid-DNA complexes might be its direct triggering of pancreatic β-cell death by activating TLR2 and TLR9, which in turn leads to the production of type I IFN and the subsequent well-defined cascade of T1D autoimmune alterations (Honda and Taniguchi 2006; Op de Beeck and Eizirik 2016). Alternatively, *E. coli*-derived amyloid might interact with antigen-presenting cells. Recent studies on other autoimmune diseases, such systemic lupus erythematosus, have shown that microbial curli fibers can lead to the activation of TLR2 and TLR9 on DCs, thus stimulating immune responses, including increased type I IFN production (Honda and Taniguchi 2006; Gallo et al., 2015; Spaulding et al., 2015; Tursi et al., 2017). Moreover, microbial amyloid has been shown to engage TLRs and trigger systemic inflammation through interaction with Peyer’s patches and lymph follicles (Teng et al., 2016; Friedland, R.P. and Chapman, M.R., 2017). We can speculate that finally, bacterial amyloid might lead to β-cell destruction and β-antigen release by stimulating IAPP aggregation and deposition in the pancreas (Westermark et al., 2017). IAPP plays multiple roles in T1D progression: on the one hand, it leads to β-cell destruction, and on the other hand, it reportedly is an autoantigen for T1D triggering (Delong et al., 2011; Denroche and Verchere, 2018). How bacterial amyloid can contribute to IAPP accumulation remains unclear; however, recent studies have revealed a trans-kingdom interplay between *E. coli* curli fibers and alpha-synuclein aggregation in the gut and brain of Parkinson’s disease models, and in triggering amyloidosis in systemic lupus erythematosus following *Salmonella typhimurium* curli-DNA treatment (Gallo et al., 2015; Chen et al., 2016).

Next, we evaluated the diabetogenic potential of *E.coli* prophages by the fact that their induction (that modulated an elevated *E.coli* phage/*E.coli* ratio from microbiome analysis) could contribute to the release of bacterial amyloid (Nanda et al., 2015). We have determined that *in vitro* activation of *E.coli* prophages with mytomicin C results in the pronounced amyloid release from preformed microbial biofilms, which according to data from metagenomics analysis, suggests that the same process occurs in the gut of those children who further develop autoimmunity and T1D (Reyrolle et al., 1982).

For *E. coli*-derived gut amyloid to activate either one of the above signaling pathways triggering autoimmune cascade and T1D, it has to reach antigen-presenting cells or pancreatic cells (Hartmann et al., 2012). This can happen in the case of increased intestinal permeability, leading to the translocation of microbial byproducts to the lamina propria, or through interaction with Peyer’s patches (Teng et al., 2016; Friedland and Chapman, 2017). Although we did not analyze intestinal permeability, altered gut barrier function is well described in T1D, with certain studies pointing it out as an important pathogenic factor for T1D (Carratu et al., 1999; Vaarala et al., 2008). Therefore, the model we propose might shed another light on the role of increased intestinal permeability–the mechanistic impact of which on T1D pathogenesis was unclear to date– suggesting that leaky gut allows *E. coli* amyloid-DNA complexes to pass to the lamina propria and trigger autoimmunity and T1D progression (Carratu et al., 1999).

We cannot conclusively explain the disappearance of *E. coli* in some control patients (who also had HLA risk for T1D); the lower abundance of *E. coli* prior to the decrease in these subjects compared to the T1D and seroconverters groups attracted our attention (Figure 4). It is possible that the lower amount of *E. coli* resulted in insufficient amyloid release to trigger autoimmunity. Another reasons for the differential effects of *E. coli* depletion across patients could be alteration of permeable state for intestinal barrier and the presence of *E. coli* strains with different levels of curli expression or amyloid synthesis under the different growth conditions; however, these possibilities need further study.

Finally, it is reasonable to speculate that the released *E. coli* phages themselves might have played a particular role in triggering T1D autoimmunity. The role of bacteriophages in the interplay with the immune system is poorly described; however, recent studies have suggested that bacterial viruses interact with eukaryotic cells, including immune cells (Górski et al., 2017 Nguyen et al., 2017).

Our analysis revealed previously overlooked correlations between an initially high level of amyloid-producing *E. coli* in the intestine, followed by their depletion, most likely due to prophage induction, and the initiation of autoimmunity and T1D progression. The diabetogenic role of *E. coli* prophages was supported by the fact that the activation of *E. coli* prophages with mitomycin C resulted in pronounced amyloid release from preformed microbial biofilms *in vitro*. Together with the data from metagenomics analysis, these findings suggested the same process might occur in the gut of children who developed autoimmunity and T1D. Combing with existing data on the immunogenic role of enterobacterial amyloid, our findings suggest that curli released by *E. coli* might trigger autoimmunity in susceptible children, highlighting the need to pay specific attention to the relationships between amyloid-producing bacteria and their bacteriophages in genetically susceptible hosts. Determining the exact role of *E. coli*-derived amyloid in the progression of T1D may lead to novel diagnostics and interventional approaches; however, obviously, further, detailed studies are required.

## Author Contributions

G.T. designed the analysis. S.B., Y.H. and G.T. conducted a metagenomics analysis. G.T. and V.T. supervised data analysis and wrote the manuscript.

## Methods

### Study Population

We used data from a prospective, longitudinal cohort study by Kostic *et al.* of 16 HLA-matched infants followed from birth until 3 years of age (Kostic *et al.*, 2016). The children, from Finland and Estonia, were recruited to the study between September 2008 and August 2010 (http://www.diabimmune.org/). The study generated shotgun metagenomics sequencing data of the fecal microbiome from 10 children who exhibited autoantibodies (six who developed serum autoantibodies with no progression to T1D, and four who were seroconverted and developed T1D) and eight non-seroconverted control individuals (Kostic *et al.*, 2016). Inclusion criteria were: presence of HLA DR-DQ alleles associated with T1D development. Data on diabetes-associated autoantibodies are presented in the original study. Demographic parameters of the study participants and data collection details are presented in Table S1.

### Microbiota Sequencing and Processing

High-throughput shotgun sequencing was performed on the Illumina HiSeq 2500 platform, generating ~2.5 Gb of sequence per sample with 101-bp paired-end reads. Human contamination was removed with the BMTagger (ftp://ftp.ncbi.nlm.nih.gov/pub/agarwala/bmtagger/). Bacterial and phage contents were quantified separately using the SRA shotgun metagenomic sequencing data. Bacterial content was quantified by taxa directly from SRA reads using Metaphalan (v. 2.0), which maps sequence reads to a database of predefined clade-specific marker genes (Segata *et al.*, 2012). All bacterial taxa with relative abundances <0.01 in all samples were excluded from statistical analysis. Phage content was assessed using a custom method. First, reads from each SRA file were assembled *de novo* into contigs with metaSPAdes (v. 3.11.1) (Nurk *et al.*, 2017). Then, contigs >200 bp were aligned to the EBI collection of phage genomes (https://www.ebi.ac.uk/genomes/phage.html) by BLAST, with a threshold e-value <1e-5 and alignment length >50% of contig length. All of the original reads were then re-mapped with Bowtie2 (v. 2.3.4.1) to the contigs with good phage BLAST matches in order to increase sensitivity and to more accurately count the abundance of reads from each type of phage (Langmead and Salzberg, 2012). Phage read counts per contig were combined per phage genome (taxa) and normalized to relative abundance. A detection threshold of two reads per sample (>90% identity to the phage genome) was used, based on a previous report (Hao *et al.*, 2018).

### Statistical Analysis of Microbial Community Composition and Differential Abundance

The QIIME pipeline (v1.9.1) was used for quality filtering of bacterial and bacteriophage DNA sequences, chimera removal (with USEARCH software), taxonomic assignment, and calculation of α-diversity, as previously described (Caporaso *et al.* 2010; Tetz *et al.*, 2017). Downstream data analysis and calculation of diversity metrics were conducted in R v3.5.1, using ggplot2 and phyloseq libraries; DESeq2 was used to calculate logarithm of fold change. Bacterial and bacteriophage communities at the genus, family, and species levels were characterized based on α- and β-diversities. α-Diversity indices (ACE, Chao 1 richness estimator, Shannon and Simpson indexes) were calculated using the phyloseq R library (McMurdie and Holmes, 2013). Differences in α-diversities between datasets were examined by the Mann-Whitney test; *p* < 0.05 was considered statistically significant.

Differences among groups where two variables exist (phenotype and time point) were analyzed by two-way ANOVA and Tukey’s multiple comparison test. When one variable was compared, an unpaired two-tailed *t*-test was used. Data were visualized using multidimensional scaling (MDS).

Correlations between *E. coli* disappearance, *Bifidobacterium* abundance, breastfeeding and antibiotic usage were assessed pairwise using the Jaccard similarity index (Jaccard, 1912). Differences were considered statistically significant at p < 0.05.

### Bacteria and Bacteriophages

*E. coli* strain MG1655 was used as a host for prophage induction experiments (Jensen 1993). Bacteria were subcultured from freezer stocks onto Mueller–Hinton agar plates (Oxoid) and incubated at 37 °C overnight. All subsequent liquid subcultures were derived from colonies isolated from these plates and were grown in Mueller–Hinton broth (Oxoid).

Bacteriophage λ from our collection was employed. Bacteriophage suspensions were routinely stored in TM buffer (10 mM Tris-HCl, 10 mM MgSO4, pH 7.2) at 4 °C. *E. coli* lysogenic strains were obtained by infection of *E. coli* MG1655 with the phage and titration of cells on Mueller–Hinton agar plates as previously described (Maniatis *et al.* 1989).

### Prophage Induction

Mitomycin C (Sigma-Aldrich) was prepared in 0.1 M phosphate buffer (pH 7.2), filtered through a 0.22-μm-pore filter (Millipore Corp., Bedford, MA), and stored at 4 °C (Georgopoulos *et al.*, 2002). A modified protocol according to Reyrolle *et al.* designed for prophage induction on 96-well plates was used (Reyrolle *et al.*, 1982; McDonald *et al.*, 2010). Twenty microliters of an overnight *E. coli* MG1655 culture, with an absorbance at 600 nm of 0.2, was dispensed in each well of a 96-well flat-bottom polystyrene tissue culture microtiter plate (Sarstedt, Numbrecht, Germany), after which 180 μl of MHB (Oxoid) was added. The plates were incubated at 37 °C for 48 h. Induction of prophages in the E. coli biofilms was provoked by the addition of mitomycin C (Sigma) to a final concentration 1 μg/ml to each well. The plates were further incubated at 37 °C and samples were taken at 4, 6, 8, and 10 h following prophage induction.

### Determination of CFU and PFU

To estimate CFU, biofilms were thoroughly scraped (Tetz *et al.*, 2009). Well contents were aspirated, placed in 1 ml of isotonic phosphate buffer (0.15 M, pH 7.2), and the total CFU number was determined by serial dilution and plating on Mueller–Hinton agar.

The number of phage virions produced after induction was estimated by phage titration (using strain MG1655 as a host) and phage plaque assay. Following induction, at the indicated times, 200-μl samples of biofilms were collected. Then, 30 μl of chloroform was added to each sample, the mixture was vortexed and centrifuged at 3000 ×*g* for 5 min in a microcentrifuge (Eppendorf 5415D). A supernatant fraction of the bacterial lysate was further used. The phage titer (number of phages per ml) was determined using the double agar overlay assay method as described previously (Yuan *et al.*, 2012). To this end, 2.5 μl of each titration point of phage lysate was spotted on Mueller−Hinton agar (Oxoid). Then, a mixture of 1 ml of *E. coli* MG1655 culture and 2 ml of 0.7% nutrient agar (heated to 45 °C) supplemented with MgSO_4_ and CaCl_2_ (to a final concentration of 10 mM each) was poured over the plate. Plates were incubated at 37 °C overnight. Each experiment was repeated in triplicate.

### Supernatant CR Depletion Assay

The amount of bacterial amyloid following prophage induction was measured in the supernatant of *E. coli* MG1655 biofilms. Biofilms were obtained as described above. Biofilm supernatant was taken at 4, 6, 8, and 10 h following prophage induction. To isolate aggregated amyloid fibers from the supernatant, the supernatant was centrifuged at 10,000 × *g*, filtered through a 0.22 µm filter to separate bacterial cells, and treated with proteinase K to eliminate contaminating proteins. CR (Sigma-Aldrich) was added to a final concentration of 10 μg/ml from a filtered stock solution of 1 mg/ml (Reichhardt *et al.*, 2015). After 5 min of equilibration, absorbance spectra were recorded from 400 to 600 nm (SmartSpec 3000, Bio-Rad). Each trace represents the average of 5 accumulated spectra. For all samples, spectra of corresponding nutrient MHB solutions with CR were used as blanks.

### Data availability

The sequencing datasets generated and/or analyzed in the current study are available from the corresponding author on reasonable request.

